# MSGPCA: Multi-Slice Graph PCA for replicate-aware Spatial Omics analysis

**DOI:** 10.64898/2026.08.05.743056

**Authors:** Anirban Chakraborty, Brian Neelon, Andrew Lawson, Peggi Angel, Dongjun Chung, Souvik Seal

## Abstract

As spatial transcriptomics (ST) and spatial proteomics (SP) technologies mature, experimental designs are increasingly moving beyond single-slice analyses toward multi-slice studies involving one or more donors and experimental conditions. Although these designs enable the identification of reproducible spatial signals, they also introduce substantial biological heterogeneity, particularly when integrating non-serial slices or anatomically distinct regions. If not modeled carefully, such variation can blur slice-specific tissue structure, mask conserved molecular patterns, and limit the discovery of biologically relevant latent structure. Although dimension reduction is essential for representing high-dimensional molecular data in a lower-dimensional space, existing multi-slice methods typically enforce a globally shared representation that inadequately accommodates slice-level heterogeneity. To address this limitation, we propose Multi-Slice Graph Principal Component Analysis (MSGPCA), which decomposes molecular variation into shared spatial factors conserved across slices and slice-specific factors that capture local tissue microarchitecture. In downstream analyses, MSGPCA-derived representations recover spatial tissue structure, denoise molecular profiles, and reveal biologically interpretable *metafeatures* associated with shared and slice-specific biology. In a mass spectrometry imaging dataset comprising nonserial slices of ductal carcinoma in situ (DCIS) and invasive breast cancer (IBC), the shared factors captured broad biological differences across tissue regions, whereas the slice-specific factors revealed intratumoral spatial variation within the IBC microenvironment. In human dorsolateral prefrontal cortex ST data, MSGPCA recovered laminar cortical architecture across adjacent slices, closely aligning with expert pathologist annotations. Together, these findings demonstrate that MSGPCA resolves shared tissue architecture while preserving local microenvironmental variation in complex multi-slice spatial omics datasets.

## 1 Introduction

Recent advances in spatial omics [1–3] enable the *in situ* profiling of diverse biomolecules within intact tissues, including genes or transcripts [4–7], lipids and metabolites [8–10], and proteins [11–14]. These technologies offer complementary views of tissue biology while varying substantially in experimental design, spatial resolution, and molecular throughput. For example, Visium HD uses a continuous lawn of 2 *µ*m barcoded squares to enable high-resolution, near-cellular whole-transcriptome mapping, whereas imaging-based platforms such as Xenium provide subcellular-resolution imaging (~0.2 *µ*m/pixel) for targeted panels of up to approximately 5,000 genes [15]. Beyond spatial transcriptomics (ST), spatial proteomics (SP) technologies such as mass spectrometry imaging (MSI) [16] can map hundreds to thousands of molecular ion features, including lipids, metabolites, peptides, and post-translational modifications, at near-cellular spatial resolutions (5–20 *µ*m). As spatial-omics technologies mature and operating costs decline, studies are increasingly shifting from single-slice analyses toward high-dimensional, multi-slice designs spanning multiple tissue slices, donors, and experimental conditions. This transition creates new opportunities to identify reproducible spatial patterns with greater potential for mechanistic validation and clinical translation [17–19]. At the same time, it introduces substantial within- and between-sample heterogeneity, which must be explicitly modeled to ensure reliable biological interpretation across analytical tasks [20].

In standard single-cell RNA sequencing (scRNA-seq) workflows and spatial-omics pipelines, including ST and MSI, dimension reduction is a crucial analytical step [21, 22]. The goal is to represent a high-dimensional feature matrix in a substantially lower-dimensional latent space, where latent factors summarize coordinated variation across molecular features. In scRNA-seq, these factors are widely used to identify cell types and cellular states; in spatial omics, they further support tasks such as spatial domain identification, feature denoising, and *metafeature* discovery, representing biologically relevant, weighted combinations of features [23–27]. Mathematically, this often involves decomposing a preprocessed molecular-abundance matrix **Y** ∈ ℝ^*n*×*p*^, rows corresponding to *n* spots or cells and columns corresponding to *p* molecular features, into a lower-dimensional factor-score matrix **Z** and a feature-loading matrix **W**, such that **Y** ≈ **ZW**^⊤^. Each latent factor is characterized by a pair of corresponding columns from **Z** and **W**. Biologically, it may represent a cell state, tissue compartment, or spatially organized molecular process. Each column of **W** defines an associated metafeature by identifying its contributing molecular features, whereas the column of **Z** shows where and how strongly that metafeature is expressed across the tissue. Classical matrix-factorization approaches include principal component analysis (PCA) [28–30], which imposes *orthogonality* constraints on the latent components or loadings, and nonnegative matrix factorization (NMF) [31, 32], which imposes *nonnegativity* constraints on the entries of **Z** and **W**. Standard PCA and NMF remain common components of spatial-omics workflows and are implemented in widely used toolboxes such as Seurat [33] and Scanpy [34]. However, because standard PCA and NMF do not explicitly model spatial dependence across spot or cell locations, substantial recent effort has focused on developing their spatially aware extensions [35–42].

Spatial extensions of PCA have long been studied in the geospatial literature [43–45]. Broadly, these methods incorporate spatial structure either by localizing PCA through geographically weighted covariance estimation [46], or by modifying the PCA objective to favor components whose scores exhibit Moran’s *I*-type spatial autocorrelation [47]. In spatial omics, this latter idea has been generalized through graph- and Gaussian process (GP)-based regularization frameworks that encourage low-dimensional representations, encoded by **Z**, to vary smoothly over tissue space. Examples include optimization-based graph-regularized methods such as GraphPCA [38], as well as probabilistic approaches such as SpatialPCA, which models latent spatial factors using GP smoothing [35]. In a similar spirit, standard NMF has been spatially augmented through probabilistic formulations such as NSF [36]. Returning to the primary focus of this manuscript, namely, multi-slice dimension reduction, several recent methods have extended these ideas to collections of *M* tissue slices or biological replicates. A common strategy is to stack or jointly model slice-level abundance matrices 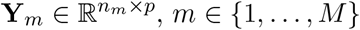, and impose a shared low-rank structure of the form **Y**_*m*_ ≈ **Z**_*m*_**W**^⊤^ where **W** denotes a shared feature-loading or metafeature matrix and **Z**_*m*_ denotes the replicate-aware low-dimensional representation for sample *m* [26, 48–51]. This formulation offers an intuitive mechanism for borrowing strength across samples, but is most naturally suited to serial, adjacent, or anatomically comparable sections, where latent factors can be interpreted as recurring tissue programs. Emerging multi-slice ST and SP studies, however, often combine non-consecutive sections, distinct anatomical regions, or heterogeneous specimens [52, 53]. In these settings, localized microarchitectures—such as immune infiltrates, stromal remodeling, or necrotic niches—may not recur across all samples [54–56]. A shared spatial representation can therefore smooth over or miss the sample-specific programs that may be most biologically informative. Although not the focus of this manuscript, neural network-based methods have also introduced graph-based nonlinear representations for single- and multi-slice ST datasets [57–61]. These approaches are effective for downstream tasks such as spatial domain identification and denoising, but their latent dimensions are generally less interpretable than linear factorization-based representations and do not naturally yield loading matrices analogous to **W**. Consequently, metafeature discovery with these methods typically relies on post hoc evaluation, whereas linear factorization-based approaches provide explicit feature loadings that more directly connect co-expressed gene modules to localized tissue microarchitectures [26, 27].

We propose multi-slice graph principal component analysis (MSGPCA), a replicate-aware framework for joint dimension reduction across multiple tissue slices. For each slice-level abundance matrix **Y**_*m*_, MSGPCA decomposes molecular variation into shared and slice-specific spatial components, 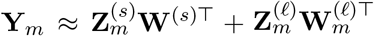, where the first term captures spatial programs conserved across slices and the second term captures localized structure unique to sample *m*. Spatial smoothness is imposed on both shared and sample-specific factors through graph-based penalties, yielding trace-regularized objectives that extend existing graph-regularized dimension-reduction methods [26, 38, 49]. The key novelty of MSGPCA is its shared-plus-specific decomposition, which separates spatial programs that are reproducible across slices from localized components that are unique to individual samples. This structure enables MSGPCA to borrow strength across replicates without forcing all tissue sections into a single common representation, thereby preserving local microarchitectures that may arise only in a subset of samples. Across simulations and real multi-slice ST and MSI datasets spanning diverse organs and disease contexts, MSGPCA yields interpretable low-dimensional representations that improve spatial domain identification, enhance feature denoising, and support biologically meaningful metafeature analysis relative to emerging alternatives.

The remainder of this paper is organized as follows. Sections 2.1 and 2.2 review GraphPCA and existing methodologies for multi-slice spatial omics analysis. In Section 2.3, we introduce MSGPCA, a unified framework for the joint analysis of multiple spatial omics sections, and discuss its downstream applications in Section 2.4. Sections 3 and 4 demonstrate the performance and practical utility of MSGPCA using simulation studies and real spatial omics datasets, respectively. We conclude with a discussion in Section 5. A scalable implementation is available as an R package on [GitHub](https://github.com/anirban0451/MSGPCA.git).

## 2 Methods

### 2.1 PCA and its graph-based spatial extensions

For a single-slice study, let **Y** ∈ ℝ^*n*×*p*^ denote the corresponding feature matrix of *p* features at *n* spatial locations. Standard PCA proceeds with decomposing **Y**’s into a product of components and loading matrices **Y** ≈ **ZW**^⊤^, such that **W** ∈ ℝ^*p*×*k*^, *k* ≪ *p*, **W**^⊤^**W** = **I**, where **I** is the identity matrix. **Z** is the collection of eigenvectors corresponding to the *k* largest eigenvalues of **W**. The orthonormality constraint on the columns of **W** is crucial as it ensures a unique decomposition. Mathematically, it can be formalized as a minimization problem of the objective function: 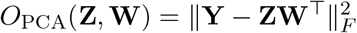, under **W**^⊤^**W** = **I**, where || · ||_*F*_ denotes the squared Frobenius norm of a matrix [62, 63]. Classical PCA, however, assumes independent observations, an assumption often violated in geostatistical settings because nearby measurements tend to be correlated, a phenomenon known as spatial autocorrelation [64]. By maximizing overall variation without considering spatial proximity, standard PCA may miss spatially coherent patterns [44, 45]. Spatially informed PCA methods address this limitation by encouraging neighboring locations to have similar principal component scores [44, 65–67]. This is particularly relevant to spatial omics, where localized tissue structures and coordinated molecular programs are of central interest [21, 68]. Motivated by this principle, Yang et al. (2024) [38] introduced an additional *L*_2_ regularization ∑_*i*~*j*_ ∥*Z*_*i*·_ − *Z*_*j*·_∥^2^ among the PC’s in **Z**, where *Z*_*i*·_ is the *i*^th^ row of **Z**, and *i* ~ *j* denotes that locations *i* and *j* are spatial neighbors. The objective function of the resulting algorithm, called GraphPCA, can be simplified as,

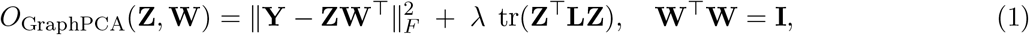

where tr(·) denotes the trace of a matrix. **L** = **D** − **A** is the unnormalized graph Laplacian, with **A** being the adjacency matrix defined as

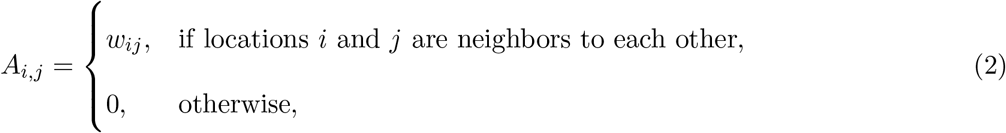

and **D** being the diagonal degree matrix given by 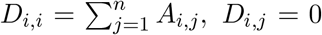 for *i*≠ *j*. For brevity, we postpone the derivation of Eq. 1 to the supplement.

In parallel to conventional PCA and its spatial variants based on penalized optimization, a substantial body of work has investigated probabilistic formulations of dimensionality reduction by placing prior distributions on latent factors and model parameters [35, 36, 40–42, 69]. These approaches naturally accommodate uncertainty quantification and have demonstrated promising performance; however, most lack a rigorous framework for jointly analyzing multiple slices.

### 2.2 Existing multi-slice graph-based PCA approaches

Now, we consider a multi-slice study comprising *M* tissue slices. For the *m*^th^ slice, let 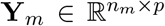n denote the feature matrix of *p* genes at *n*_*m*_ spatial locations, *m* = 1, 2, …, *M*. One possible dimension-reduction strategy is to apply GraphPCA separately to each slice, yielding the decomposition 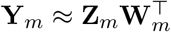 by minimizing the slice-specific version of the objective function in Eq. 1. However, because the model parameters are estimated independently for each slice, this approach makes it difficult to identify reproducible spatial programs shared across slices. Consequently, downstream analyses such as global domain detection and metafeature discovery require post hoc alignment of the latent factors, a process that can become increasingly cumbersome as the number of slices grows. Several methods instead impose a shared loading matrix, **W**_*m*_ = **W** for all *m* ∈ 1, …, *M* [26, 49], and jointly estimate the parameters using a combined objective function:

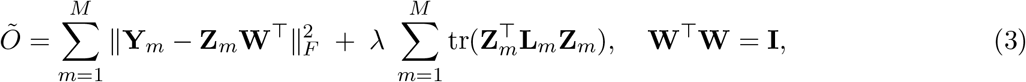

that results in a closed-form solution.

Although such a formulation is well suited to jointly analyzing serial tissue sections [26], multi-slice studies may instead comprise nonconsecutive sections sampled from different regions within the same tissue specimen or from distinct specimens altogether (Figure 4a). In such settings, individual slices may capture region-specific tumor microenvironments and spatial programs that are not consistently represented across the remaining slices. The shared formulation in Eq. 3 may adequately identify global spatial domains, but it cannot explicitly distinguish these localized, slice-specific programs from patterns shared across slices. This limitation motivates the development of our replication-aware model that simultaneously captures globally shared spatial architectures and slice-specific heterogeneity.

### 2.3 Proposed method: MSGPCA

We decompose the slice-specific gene expression matrices into *k*_*s*_ shared components and *k*_*ℓ*_ slice-specific components,

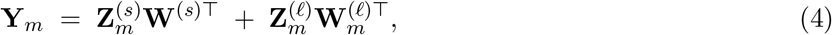

where 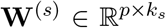 and 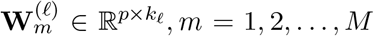 are orthonormal matrices denoting the shared and slice-specific loadings, and 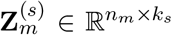 and 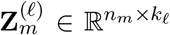 denote shared and slice specific PCs, respectively. Note that when 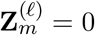, we get back a decomposition similar to Eq. 3.

Unlike previous approaches [26, 38], imposing orthonormality on the shared loading matrix **W**^(*s*)^ and slice-specific loading matrix 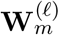 is no longer sufficient to ensure a unique solution (see Supplementary Material). To address this identifiability issue, we additionally require each 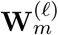 to be orthogonal to **W**^(*s*)^,

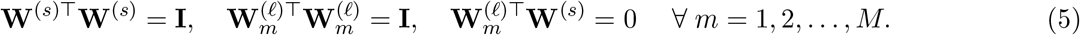

Next, we enforce spatial smoothness in both the shared and slice-specific PC scores, 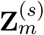 and 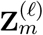, using trace penalties analogous to that in Eq. (3),

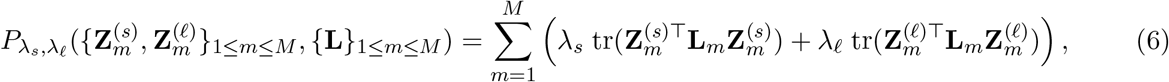

where **L**_*m*_ = **A**_*m*_ − **D**_*m*_ denotes the *m*^th^ graph Laplacian. Finally, the penalized objective function is

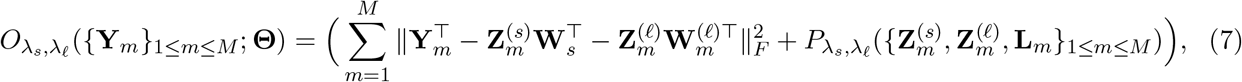

which is mimized w.r.t. the full set of parameters, 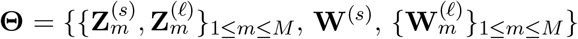. We use an iterative optimization procedure that alternately updates the model parameters and evaluates the objective function. The algorithm terminates when the relative change in the objective function falls below a prespecified convergence threshold. The complete procedure is summarized in Algorithm 1, with detailed derivations provided in the Supplementary Material.

The performance of Algorithm 1 depends on the choice of spatial adjacency graphs, represented by 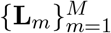, and the smoothing parameters *λ*_*s*_ and *λ*_*ℓ*_. In practice, we construct *k*-nearest-neighbor (*k*-NN) graphs with *k* ≤ 6, as larger values produced excessive smoothing of the estimated factors in our empirical experiments. On the other hand, a large value for *λ*_*ℓ*_ (and *λ*_*s*_) fails to capture slice-specific variation, while a small *λ*_*ℓ*_ (and *λ*_*s*_) introduces artifacts in the factors that obscure true biological signals. We utilize a Pareto front-based optimization framework to select the best combination of *λ*_*s*_ and *λ*_*ℓ*_ [70, 71]. The estimated parameters 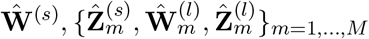 facilitate several downstream applications as listed in the following section.

#### Algorithm 1

MSGPCA: complete algorithm

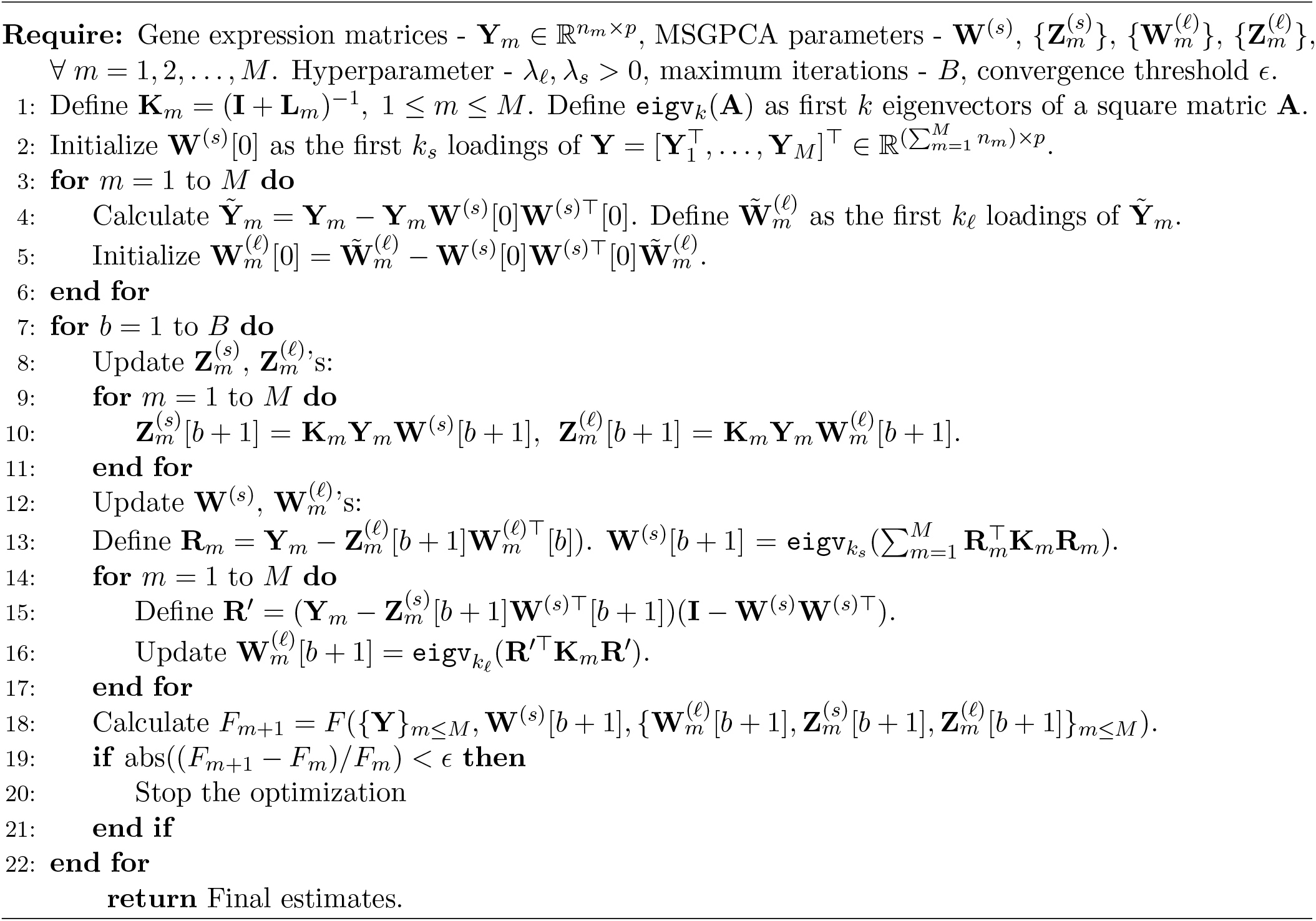

### 2.4 Downstream applications

The resulting PCs support several important downstream applications, including spatial domain detection, whereby cells or spots are grouped into clusters based on similarities in their joint molecular profiles [57]; feature-expression denoising, whereby technical noise is reduced by reconstructing the data from a lower-dimensional representation [72]; and the discovery of shared and slice-specific metafeatures, defined as weighted combinations of molecular features that capture coordinated variation across cells or spots [26].

- **Spatial domain detection**. To enable joint domain detection across all tissue sections, we vertically concatenate the estimated shared factor-score matrices, 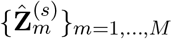, to form a global representation of dimension 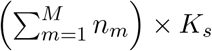, where *K*_*s*_ denotes the number of shared factors. We then apply probabilistic clustering to the aggregated representation using a Gaussian mixture model (GMM) implemented in the *mclust* R package [58, 73, 74], followed by spatial refinement to correct isolated or spatially inconsistent domain assignments [35].
- **Feature expression denoising**. Using the PCs, the denoised feature expression for the *m*^th^ slice can be reconstructed as: 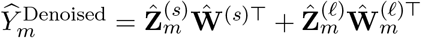. By retaining the dominant molecular variation captured by the shared and slice-specific factors while reducing residual measurement noise, this low-rank reconstruction can enhance spatially coherent expression patterns [72, 75].
- **Metafeature discovery**. Each column of **Ŵ** ^(*s*)^ defines a shared metafeature, represented as a weighted combination of the original molecular features, while the corresponding columns of 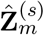 characterize its spatial activity within each slice. These shared metafeatures capture reproducible molecular patterns across tissue sections. Zhang et al. [26] used a related approach to identify anatomy-preserving metagenes in ST; however, their formulation does not explicitly model molecular patterns that are unique to individual slices, which may be important when the sampled regions are biologically heterogeneous. In contrast, the slice-specific components of MSGPCA, 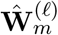 and 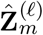, capture molecular signatures and their spatial distributions that are not explained by the shared component. Specifically, each column of 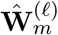 defines a slice-specific metafeature, and the corresponding column of 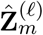 describes its spatial activity. This decomposition enables replication-aware analysis of both reproducible molecular structure and region-specific tissue heterogeneity across multiple slices.

## 3 Simulation experiment

To demonstrate the utility of MSGPCA in decoupling global tissue architecture from localized spatial heterogeneity, we adapted the “ggblocks” simulation framework from Townes et al. (2023)[36]. Following Wang et al. (2025) [50], we defined four binary spatial domains across a uniform coordinate grid (Figure 1a). We simulated two distinct tissue slices and generated factor-preserving genes, designing them to exhibit high expression strictly when their corresponding spatial factor was active. To accurately reflect the high overdispersion and dropout rates characteristic of ST, we simulated these gene counts using a mixture of Bernoulli and Negative Binomial distributions. Because heterogeneous biological samples frequently contain regions with structurally mismatched expression boundaries, we introduced targeted spatial perturbations into a subset of features exclusively within the second slice (Figure 1b, bottom panel). We intentionally left the corresponding baseline expression in the first slice unaltered. Next, we integrated non-spatially structured background genes following [36] and [50] to emulate realistic baseline noise profiles. Finally, we generated *n* = 50 independent simulation replicates, configuring each slice to contain *p* = 500 genes. To benchmark our algorithm for spatial domain identification, we used two methods, STAGATE and GraphST, that showed satisfactory performance in a recent domain detection benchmark study [76]. We also selected jsPCA, a recently developed spatial domain-detection method for multi-slice data [74]. Details of these methods are summarised in Table 1.

**Table 1.** Details of competing methods used for benchmarking. Among these listed methods, STAGATE requires alignment for multi-slice data analysis.

| Method | Algorithm | Spatial Clustering | Denoising | Reproducible spatial programs | Slice-specific spatial programs |
| --- | --- | --- | --- | --- | --- |
| MSGPCA | Replicate-aware spatially regularized PCA with slice-specific components | ✓ | ✓ | ✓ | ✓ |
| jsPCA[74] | Joint PCA on spatially smoothened covariance matrix | ✓ | ✓ | ✓ | × |
| STAGATE[59] | Graph-attention autoencoder | ✓ | ✓ | × | × |
| GraphST[58] | Graph-based self-supervised contrastive learning | ✓ | × | × | × |

**Figure 1.**
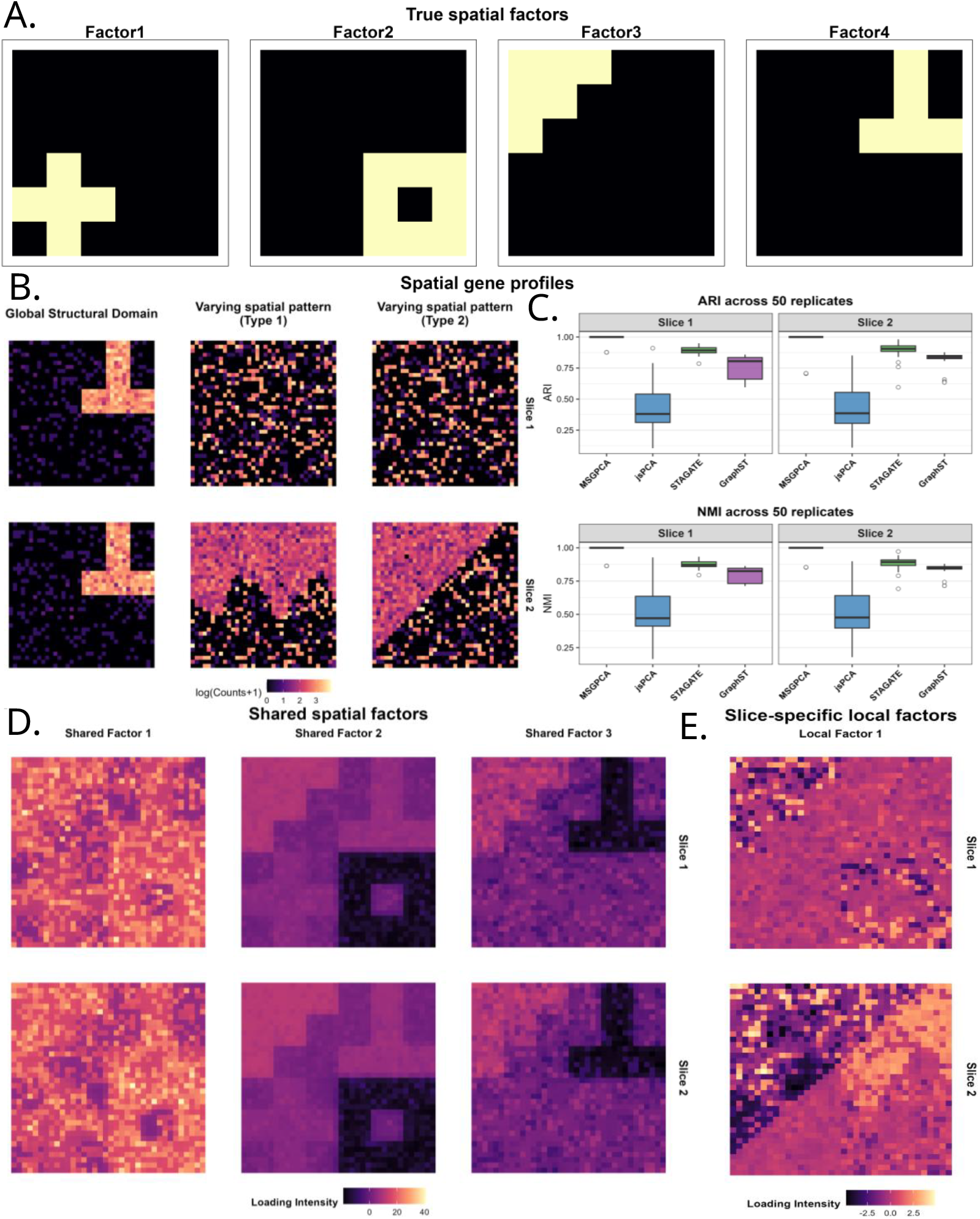
(A). True spatial distributions of the factors used in the simulation study. For each of the four factors, yellow indicates presence (factor value of 1), whereas black indicates absence (factor value of 0) at the corresponding spatial location. (B). Genes of different spatial patterns across two slices. Global genes that preserve the factors from (A) in both Slices 1 and 2. Type 1 and 2 genes that are completely noisy in Slice 1, but have spatial expression in Slice 2. (C). Boxplot of ARI and NMI values for four methods across 50 simulated datasets. (D). First 3 shared factors for 2 slices in a specific dataset simulated using (A). (E). First local factors for two slices corresponding to (D).

Over 50 replicated datasets, MSGPCA achieved exceptional clustering accuracy across all replicates (median Adjusted Rand Index, ARI = 1.00), significantly outperforming competing methods including jsPCA (median ARI = 0.38), GraphST (median ARI = 0.82), and STAGATE (median ARI = 0.90) (Figure 1c). Moreover, our shared factors successfully captured the conserved global spatial patterns, cleanly reconstructing the fundamental structural boundaries (Figure 1d; see Supplementary Material). The first factor mainly captured the background noise, and could be interpreted as an embedding that facilitated capturing the unimportant features. We could safely discard such embedding in downstream analyses. Crucially, our slice-specific local factors successfully isolated the precise diagonal and sinusoidal perturbations that we injected into slice 2 (Figure 1e; see Supplementary Material). For slice 1, which lacked targeted spatial anomalies, the local factors appropriately modeled only residual stochastic noise. This ultimately confirms the capacity of MSGPCA to strictly partition highly specific localized spatial anomalies from conserved global structures.

## 4 Real data analysis

### 4.1 Human dorsolateral prefrontal cortex Visium data

We analyzed a human dorsolateral prefrontal cortex (DLPFC) Visium dataset comprising twelve tissue sections from three donors [77]. Each section contained a median of 3,844 spots and 33,538 measured genes. Our analysis focused on four serial sections from Donor 3 (Br5595), each of which included five manually annotated regions: four cortical layers and white matter. We applied minimal spot-level filtering and retained the top 4,000 highly variable genes for analysis. We next applied MSGPCA with *k*_*s*_ = 15 shared factors and *k*_*l*_ = 5 local factors, and performed downstream analyses detailed in Section 2.4.

We used NMI to compare the domain detection performance of the four methods. MSGPCA consistently uncovered the true spatial domains across all four slices (Figure 2A). MSGPCA achieved a mean NMI of 0.82, substantially outperforming jsPCA (0.36), STAGATE (0.40), and GraphST (0.32). Next, to demonstrate MSGPCA’s utility in marker denoising, we examined the normalized and MSGPCA-estimated expressions of five established layer-specific markers [77] in slice 151669: CARTPT (Layer 3), RORB (Layer 4), PCP4 (Layer 5), KRT17 (Layer 6), and MOBP (white matter). Among these markers, RORB exhibited particularly noisy expression patterns, likely due to the narrow anatomical extent of Layer 4. MSGPCA substantially reduced this technical noise while preserving the expected layer-specific spatial localization, thereby recovering a clearer underlying expression pattern (Figure 2B). Furthermore, MSGPCA consistently increased the median expression levels of these marker genes across tissue slices, yielding improved signal recovery and clear laminar structure (Figures 2C, D). Lastly, our method recovered reproducible spatial programs across all four slices that reflect the conserved laminar architecture of the cortex, with distinct factors highlighting mid-cortical laminae and deep cortical regions near the white matter boundary (Figure 2E).

**Figure 2.**
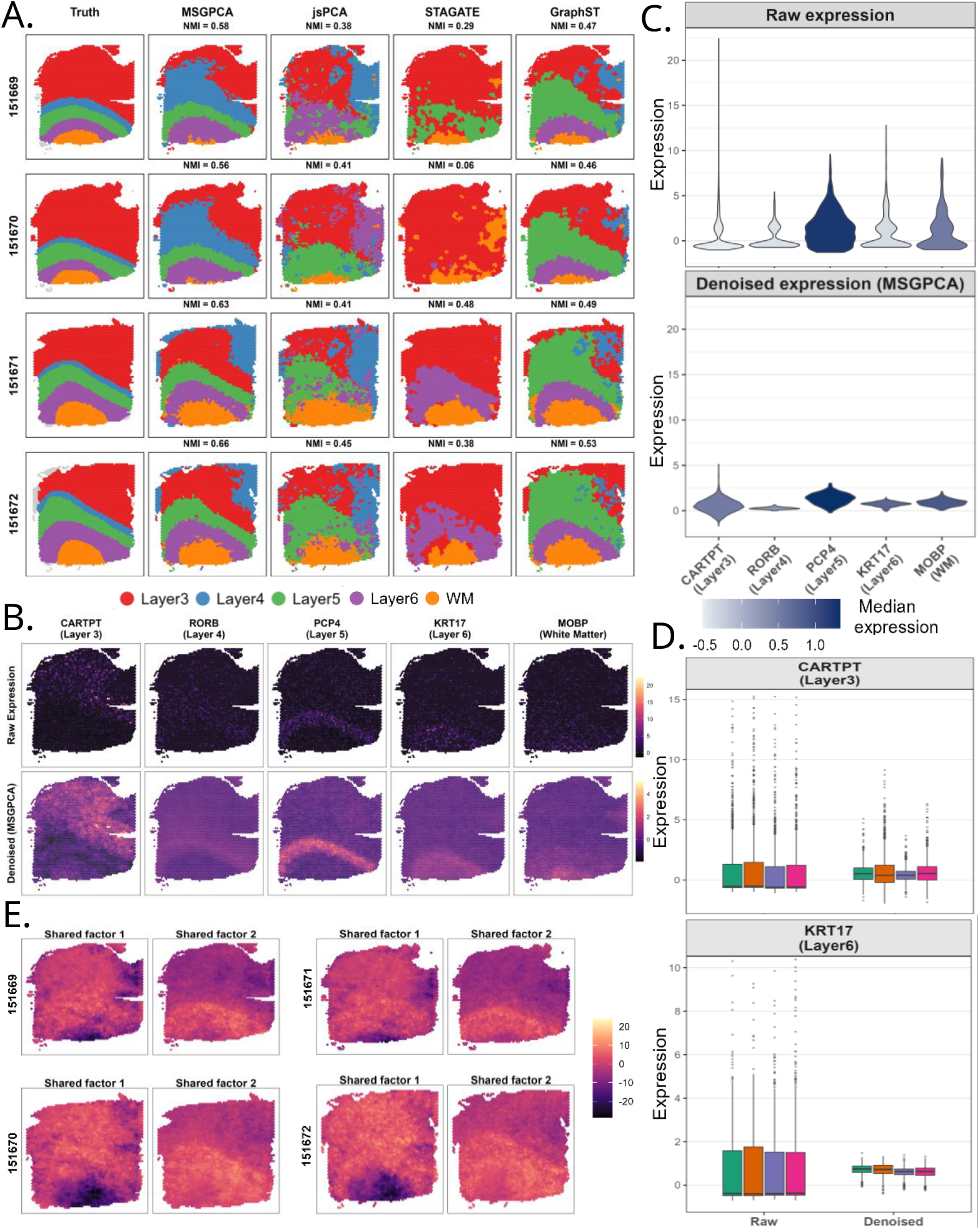
(A). Clustering results for four methods in DLPFC slices 151669, 151670, 151671 and 151672. The corresponding NMI values are provided in parentheses. (B). Raw and denoised expression of 5 layer-specific marker genes in slice 151671. (C). Violin plot of raw and denoised expression of the same markers. (D). Boxplot of the raw and denoised gene expressions for markers CARTPT (layer 3) and KRT17 (layer 5) across slices 151669 – 151672. The colors are indicators of different slices. (E). First two shared factors for all the 4 slices. Across all plots, *raw* expression refers to the final values after routine preprocessing and filtering.

### 4.2 Hypothalamus MERFISH data

Next, we analyzed an ST dataset comprising five adjacent tissue slices from the hypothalamic preoptic region, profiled via MERFISH [78]. Each slice captured expression profiles for a conserved set of 155 targeted genes with eight annotated anatomical domains: the third ventricle (V3), bed nuclei of the stria terminalis (BST), columns of the fornix (fx), medial preoptic area (MPA), medial preoptic nucleus (MPN), periventricular hypothalamic nucleus (PV), paraventricular hypothalamic nucleus (PVH), and paraventricular nucleus of the thalamus (PVT). Following global mean-variance scaling of the expression matrices across all slices, we applied MSGPCA with *k*_*s*_ = 15 shared factors and *k*_*l*_ = 5 local factors, and performed downstream analyses detailed in Section 2.4. MSGPCA achieved superior clustering performance across all five slices relative to baseline methods (Figure 3A). Furthermore, we performed UMAP with the first four latent factors of MSGPCA that revealed two structural domains, V3 and BST, separated from the remaining clusters (Figure 3B). In contrast, GraphST and STAGATE achieved only marginal separation for V3, whereas jsPCA failed to detect any separation. These findings indicate that the low-dimensional representations generated by MSGPCA encapsulate spatial variances that are highly predictive of true anatomical architecture. Notably, none of the methods accurately identified the PVA or MPN, underscoring the difficulty of resolving these highly localized subtissue structures.

**Figure 3.**
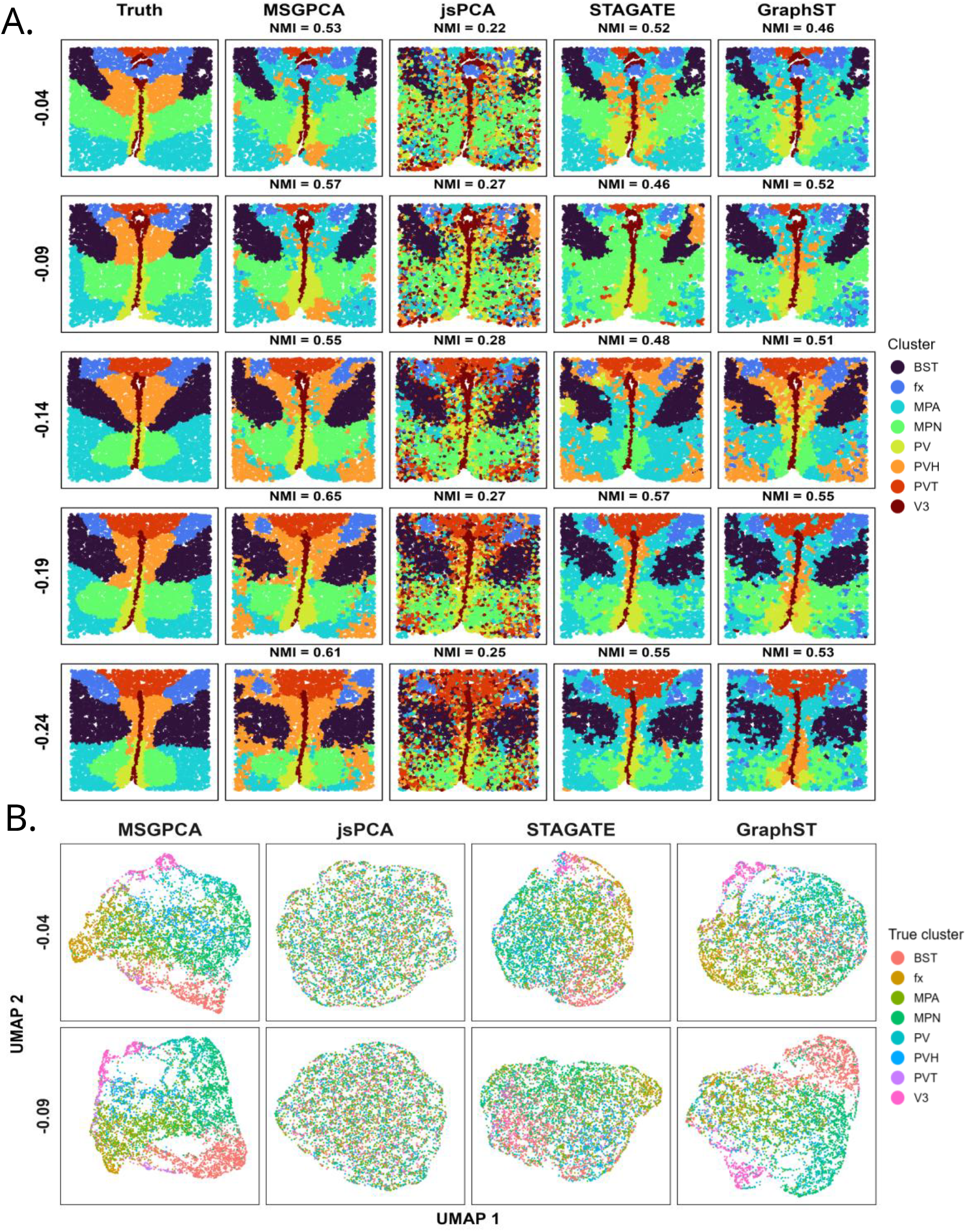
(A). Clustering results for four methods across the 5 preoptic region slices from Moffit et al. (2018) [78]. (B). UMAP based on the factors obtained using different methods in two representative slices, colored by true cluster labels.

### 4.3 DCIS and IBC MALDI data

As discussed previously, sample-specific spatial programs may be particularly pronounced in nonserial tissue sections sampled from distinct regions of the same specimen. To illustrate this setting, we applied MSGPCA to a multi-slice MALDI-MSI spatial proteomics dataset comprising multiple pathologist-annotated ductal carcinoma in situ (DCIS), invasive breast cancer (IBC), and normal breast tissue slices from an ongoing MUSC study investigating the spatial distributions of collagen peptides and immune cells (Figure 4A). Pathologically, DCIS—which constitutes 20% of incident breast cancer cases—is defined by the localized expansion of malignant epithelial cells that remain strictly compartmentalized within the milk ducts without breaching the adjacent stroma [79]. Conversely, IBC is an aggressive invasive phenotype driven by severe lymphovascular infiltration, wherein malignant emboli obstruct the dermal lymphatic channels [80]. Up to half of untreated patients with DCIS develop IBC over a 10-year period [81]. Consequently, current medical consensus mandates surgical excision for all patients presenting with DCIS.

**Figure 4.**
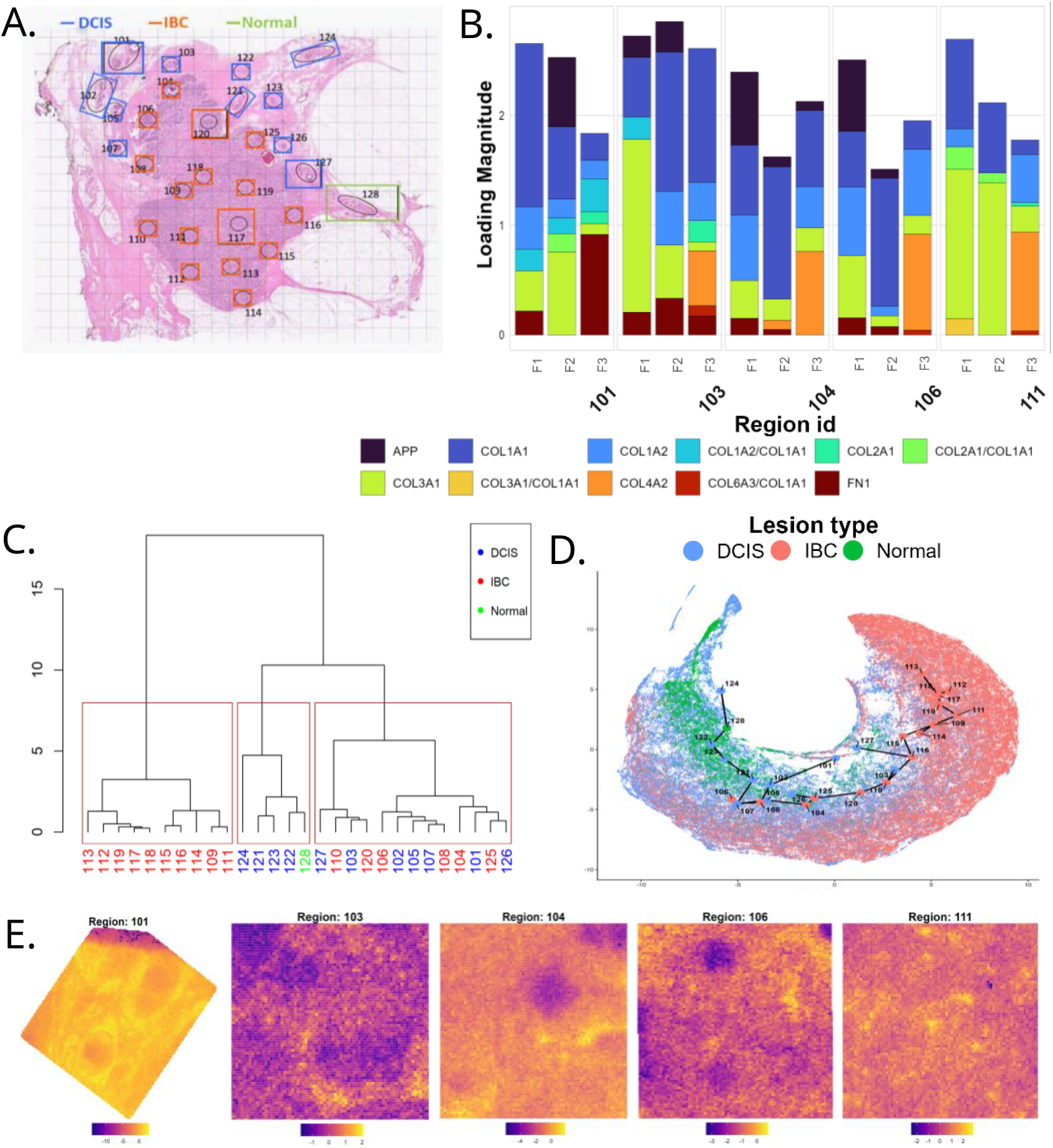
(A). H&E picture of the breast tissue section which annotated section. (B). Weight distribution of different proteins in the first three factors across five selected regions - 101, 103, 104, 106 and 114, plotted as a stacked bar. The colors denote specific proteins. (C). Hierarchical clustering based on region-specific mean shared factors. For each region, these mean shared factors are calculated by averaging over all the spots. Red boxes highlight unsupervised clustering result with three clusters. (D). UMAP of all regions with corresponding shared factors. The large points denote the region specific centroids. The lines connecting them summarize a global relationship between these regions. (E). Raw expression of the regions corresponding to b. Similar to Figure 2, here *raw* expression refers to original expression after necessary preprocessing and scaling.

The complete dataset comprised 11 DCIS regions of interest (ROIs), 16 IBC ROIs, and 1 normal tissue ROI, with a total of 199,200 pixels with 56 features. After filtering excessively sparse features and performing feature-wise intensity scaling across all slices, we retained 39 features. We subsequently applied MSGPCA to this preprocessed dataset, specifying *k*_*s*_ = 5 shared and *k*_*l*_ = 3 slice-specific latent factors. We next computed the shared-factor profiles for each ROI by averaging across its constituent pixels and performed hierarchical clustering to group the ROIs. Several IBC and DCIS regions consistently co-clustered, suggesting the presence of shared molecular programs across histologically distinct disease states. This observation was further supported by UMAP visualization of the shared factors. Although most regions followed the expected continuum from DCIS to IBC, several deviated from this trajectory and formed distinct branches (Figure 4C, D). Together, these findings indicate that some slices may have unique molecular activities which is not completely captured by the shared factors. Therefore, to examine slice-specific spatial programs, we selected a representative subset of two DCIS and three IBC sections.

Among the identified metafeatures, regions 101, 104, and 106 demonstrated strong signals for the amyloid precursor protein (APP). In contrast, APP presence was notably low in region 103 and entirely absent from the spatial programs of region 111 (Figure 4B). spatial autocorrelation in regions 101 and 104 (Figure 4E, Moran’s *I* = 0.88, 0.82 respectively), characterized by distinct, localized clusters of high and low intensity. Region 106 demonstrated a comparatively moderate spatial correlation (Moran’s *I* = 0.65), driven primarily by localized zones of low expression, while regions 103 and 111 lacked meaningful spatial structure (Moran’s *I* = 0.55 and 0.48, respectively). Elevated APP is a documented feature of IBC tumors [82]. Because breast carcinomas exhibit profound patient- and tissue-level variability [83], the ability of MSGPCA to capture these varying APP expression patterns validates its utility. By isolating tumor-specific metafeatures after subtracting spatially reproducible global programs (see the Supplementary Material), MSGPCA provides a highly effective mechanism for elucidating intra-tumor heterogeneity. Moreover, by delineating aggressive subregions among the non-invasive DCIS lesion, MSGPCA highlights spatially emergent malignant programs, thereby underscoring its value for early detection of high-risk disease.

## 5 Discussion

As spatial omics technologies mature and become more affordable, studies profiling multiple tissue slices are becoming increasingly common. Such datasets can exhibit substantial biological variation within and across samples, particularly when slices are collected from nonadjacent or anatomically distinct regions. Although existing methods are generally well suited to serial or closely related slices, they may inadequately represent heterogeneity across more distinct tissue regions. To address this limitation, we developed MSGPCA, a joint dimensionality-reduction framework that decomposes each feature matrix into a shared component that captures spatially organized molecular patterns reproducible across slices and a slice-specific component that captures patterns unique to each tissue slice. We developed an efficient, user-friendly R package implementing MSGPCA, which is available on [GitHub](https://github.com/anirban0451/MSGPCA.git).

Across extensive simulation studies, MSGPCA achieved superior spatial domain detection performance in comparison to popular existing methods. In analyses of the human DLPFC dataset, MSGPCA more consistently resolved the cortical layers across all four sections and, through expression denoising, enhanced the median expression of layer-specific marker genes within their corresponding regions. Its shared components also preserved the reproducible laminar architecture across sections. In the mouse hypothalamus dataset, MSGPCA showed improved clustering performance for several preoptic regions. Finally, in the MALDI-MSI proteomics dataset, MSGPCA identified slice-specific metafeatures distinguishing heterogeneous DCIS and IBC regions, including spatially variable APP expression. Together, these downstream analyses demonstrate the ability of MSGPCA to capture both reproducible tissue organization and region-specific molecular heterogeneity across nonadjacent or biologically distinct sections.

A potential limitation of MSGPCA is that both shared and slice-specific factors may capture localized spatial noise, producing redundant or weakly informative metafeatures that require post hoc filtering before biological interpretation. Future work, therefore, could incorporate sparsity directly into the loading matrices, for example through an additional LASSO penalty [26], to yield more compact and interpretable molecular signatures. The current framework also requires users to specify the number of factors, which is often selected heuristically; probabilistic spatial factor models with shrinkage priors could instead adaptively determine the effective dimensionality [84]. MSGPCA could further be extended to distinguish donor- and condition-specific variation in multi-donor, multi-slice studies [48, 85]. In addition, GMM-based domain detection became less stable as the number of latent factors increased, reflecting the broader challenges of high-dimensional clustering [86, 87]. Integrating regularized, spatially informed clustering methods may improve robustness in such settings [23, 88–90]. Finally, future Bayesian extensions could replace the Gaussian observation model with data-type-specific likelihoods, such as negative binomial distributions for sequencing counts and flexible skewed continuous distributions for continuous peptide intensities [91, 92].

