## Supplementary Material for "MSGPCA: Multi-Slice Graph PCA for replicate-aware Spatial Omics analysis"


#### S1 Derivation of GraphPCA objective function

The GraphPCA penalty [1] encourages neighboring locations to have similar low-dimensional representations through the term

$$\sum_{i\sim j} \|Z_{i\cdot} - Z_{j\cdot}\|^2.$$

Let $\mathbf{A}$ denote the adjacency matrix of the spatial graph, with $A_{ij} = 1$ if locations $i$ and $j$ are neighbors and $A_{ij} = 0$ otherwise. Defining the degree matrix $\mathbf{D} = \text{diag}(d_1, \ldots, d_n)$, where $d_i = \sum_j A_{ij}$, and the graph Laplacian $\mathbf{L} = \mathbf{D} - \mathbf{A}$, the penalty can be rewritten as

$$\sum_{i\sim j} \|Z_{i\cdot} - Z_{j\cdot}\|^2 = \sum_{i,j} A_{ij}\|Z_{i\cdot} - Z_{j\cdot}\|^2.$$

Expanding the squared norm yields

$$\sum_{i,j} A_{ij} \left( Z_{i\cdot}^\top Z_{i\cdot} + Z_{j\cdot}^\top Z_{j\cdot} - 2 Z_{i\cdot}^\top Z_{j\cdot} \right),$$

which simplifies to

$$2\,\mathrm{tr}(\mathbf{Z}^\top \mathbf{D}\mathbf{Z}) - 2\,\mathrm{tr}(\mathbf{Z}^\top \mathbf{A}\mathbf{Z}) = 2\,\mathrm{tr}(\mathbf{Z}^\top \mathbf{L}\mathbf{Z}).$$

Since the multiplicative constant can be absorbed into the tuning parameter $\lambda$, the neighborhood smoothness penalty is equivalently expressed as

$$\mathrm{tr}(\mathbf{Z}^\top \mathbf{L}\mathbf{Z}),$$

leading to the GraphPCA objective in Equation (1).

### S2 Orthonormality conditions in MSGPCA

The orthonormality constraints on $\mathbf{W}^{(s)}$ and $\mathbf{W}_m^{(\ell)}$ alone do not guarantee identifiability in Eq. (4). To see this, consider an arbitrary matrix $\mathbf{R} \in \mathbb{R}^{k_\ell \times k_s}$ and define

$$\widetilde{\mathbf{Z}}_m^{(s)} = \mathbf{Z}_m^{(s)} - \mathbf{Z}_m^{(\ell)}\mathbf{R}, \qquad \widetilde{\mathbf{W}}_m^{(\ell)\top} = \mathbf{W}_m^{(\ell)\top} + \mathbf{R}\mathbf{W}^{(s)\top}.$$

Then,

$$\begin{aligned}\widetilde{\mathbf{Z}}_m^{(s)}\mathbf{W}^{(s)\top} + \mathbf{Z}_m^{(\ell)}\widetilde{\mathbf{W}}_m^{(\ell)\top} &= \left(\mathbf{Z}_m^{(s)} - \mathbf{Z}_m^{(\ell)}\mathbf{R}\right)\mathbf{W}^{(s)\top} + \mathbf{Z}_m^{(\ell)}\left(\mathbf{W}_m^{(\ell)\top} + \mathbf{R}\mathbf{W}^{(s)\top}\right) \\ &= \mathbf{Z}_m^{(s)}\mathbf{W}^{(s)\top} + \mathbf{Z}_m^{(\ell)}\mathbf{W}_m^{(\ell)\top}.\end{aligned}$$

Hence, infinitely many pairs $\left(\widetilde{\mathbf{Z}}_m^{(s)}, \widetilde{\mathbf{W}}_m^{(\ell)}\right)$ produce exactly the same reconstruction of $\mathbf{Y}_m$. This ambiguity arises because variation can be transferred between the shared and slice-specific components without changing the fitted value. Consequently, the decomposition is not unique when only the individual orthonormality conditions $\mathbf{W}^{(s)\top}\mathbf{W}^{(s)} = \mathbf{I}$ and $\mathbf{W}_m^{(\ell)\top}\mathbf{W}_m^{(\ell)} = \mathbf{I}$ are imposed. Requiring the additional cross-orthogonality constraint $\mathbf{W}_m^{(\ell)\top}\mathbf{W}^{(s)} = \mathbf{0}$ eliminates this overlap and separates the shared and slice-specific loading spaces.

#### S3 Derivation of the MSGPCA Updates

We derive the alternating updates for the MSGPCA objective. For simplicity, write

$$\mathbf{Y}_m = \mathbf{Z}_m^{(s)}\mathbf{W}^{(s)\top} + \mathbf{Z}_m^{(\ell)}\mathbf{W}_m^{(\ell)\top} + \mathbf{E}_m,$$

where

$$\mathbf{W}^{(s)\top}\mathbf{W}^{(s)} = \mathbf{I}, \qquad \mathbf{W}_m^{(\ell)\top}\mathbf{W}_m^{(\ell)} = \mathbf{I}, \qquad \mathbf{W}_m^{(\ell)\top}\mathbf{W}^{(s)} = \mathbf{0}.$$

The penalized objective is

$$\begin{aligned} O_{\lambda_s,\lambda_\ell} = &\sum_{m=1}^{M} \left\| \mathbf{Y}_m - \mathbf{Z}_m^{(s)}\mathbf{W}^{(s)\top} - \mathbf{Z}_m^{(\ell)}\mathbf{W}_m^{(\ell)\top} \right\|_F^2 \\ &+ \sum_{m=1}^{M} \left\{ \lambda_s \operatorname{tr}\left(\mathbf{Z}_m^{(s)\top}\mathbf{L}_m\mathbf{Z}_m^{(s)}\right) + \lambda_\ell \operatorname{tr}\left(\mathbf{Z}_m^{(\ell)\top}\mathbf{L}_m\mathbf{Z}_m^{(\ell)}\right) \right\}. \end{aligned} \tag{1}$$

**Update for $\mathbf{Z}_m^{(s)}$.** Fixing $\mathbf{W}^{(s)}$, $\mathbf{W}_m^{(\ell)}$, and $\mathbf{Z}_m^{(\ell)}$, the part of the objective involving $\mathbf{Z}_m^{(s)}$ is

$$\left\| \mathbf{Y}_m - \mathbf{Z}_m^{(s)}\mathbf{W}^{(s)\top} - \mathbf{Z}_m^{(\ell)}\mathbf{W}_m^{(\ell)\top} \right\|_F^2 + \lambda_s \operatorname{tr}\left(\mathbf{Z}_m^{(s)\top}\mathbf{L}_m\mathbf{Z}_m^{(s)}\right).$$

Taking the derivative with respect to $\mathbf{Z}_m^{(s)}$ and setting it equal to zero gives

$$-2\left(\mathbf{Y}_m - \mathbf{Z}_m^{(s)}\mathbf{W}^{(s)\top} - \mathbf{Z}_m^{(\ell)}\mathbf{W}_m^{(\ell)\top}\right)\mathbf{W}^{(s)} + 2\lambda_s\mathbf{L}_m\mathbf{Z}_m^{(s)} = \mathbf{0}.$$

Using

$$\mathbf{W}^{(s)\top}\mathbf{W}^{(s)} = \mathbf{I}, \qquad \mathbf{W}_m^{(\ell)\top}\mathbf{W}^{(s)} = \mathbf{0},$$

we obtain

$$-\mathbf{Y}_m\mathbf{W}^{(s)} + \mathbf{Z}_m^{(s)} + \lambda_s\mathbf{L}_m\mathbf{Z}_m^{(s)} = \mathbf{0}.$$

Therefore,

$$\left(\mathbf{I}_{n_m} + \lambda_s\mathbf{L}_m\right)\mathbf{Z}_m^{(s)} = \mathbf{Y}_m\mathbf{W}^{(s)}.$$

Thus the update is

$$\boxed{\mathbf{Z}_m^{(s)} = \mathbf{K}_{m,s}\mathbf{Y}_m\mathbf{W}^{(s)}, \qquad \mathbf{K}_{m,s} = \left(\mathbf{I}_{n_m} + \lambda_s\mathbf{L}_m\right)^{-1}.} \tag{2}$$

**Update for $\mathbf{Z}_m^{(\ell)}$.** Similarly, fixing $\mathbf{W}^{(s)}$, $\mathbf{W}_m^{(\ell)}$, and $\mathbf{Z}_m^{(s)}$, the derivative with respect to $\mathbf{Z}_m^{(\ell)}$ gives

$$-2\left(\mathbf{Y}_m - \mathbf{Z}_m^{(s)}\mathbf{W}^{(s)\top} - \mathbf{Z}_m^{(\ell)}\mathbf{W}_m^{(\ell)\top}\right)\mathbf{W}_m^{(\ell)} + 2\lambda_\ell\mathbf{L}_m\mathbf{Z}_m^{(\ell)} = \mathbf{0}.$$

Using

$$\mathbf{W}_m^{(\ell)\top}\mathbf{W}_m^{(\ell)} = \mathbf{I}, \qquad \mathbf{W}^{(s)\top}\mathbf{W}_m^{(\ell)} = \mathbf{0},$$

we obtain

$$\left(\mathbf{I}_{n_m} + \lambda_\ell\mathbf{L}_m\right)\mathbf{Z}_m^{(\ell)} = \mathbf{Y}_m\mathbf{W}_m^{(\ell)}.$$

Hence,

$$\boxed{\mathbf{Z}_m^{(\ell)} = \mathbf{K}_{m,\ell}\mathbf{Y}_m\mathbf{W}_m^{(\ell)}, \qquad \mathbf{K}_{m,\ell} = \left(\mathbf{I}_{n_m} + \lambda_\ell\mathbf{L}_m\right)^{-1}.} \tag{3}$$

**Update for $\mathbf{W}^{(s)}$.** To update the shared loading matrix, fix the slice-specific terms and define the partial residual

$$\mathbf{R}_m^{(s)} = \mathbf{Y}_m - \mathbf{Z}_m^{(\ell)}\mathbf{W}_m^{(\ell)\top}.$$

Then the shared part of the objective becomes

$$\sum_{m=1}^{M}\left\|\mathbf{R}_m^{(s)} - \mathbf{Z}_m^{(s)}\mathbf{W}^{(s)\top}\right\|_F^2 + \lambda_s\sum_{m=1}^{M}\operatorname{tr}\left(\mathbf{Z}_m^{(s)\top}\mathbf{L}_m\mathbf{Z}_m^{(s)}\right).$$

For fixed $\mathbf{W}^{(s)}$, the optimal score matrix is

$$\mathbf{Z}_m^{(s)} = \mathbf{K}_{m,s}\mathbf{R}_m^{(s)}\mathbf{W}^{(s)}, \qquad \mathbf{K}_{m,s} = (\mathbf{I}_{n_m} + \lambda_s \mathbf{L}_m)^{-1}.$$

Substituting this expression into the objective and removing terms that do not depend on $\mathbf{W}^{(s)}$ shows that minimizing the objective is equivalent to maximizing

$$\operatorname{tr}\left[\mathbf{W}^{(s)\top}\left\{\sum_{m=1}^{M}\mathbf{R}_m^{(s)\top}\mathbf{K}_{m,s}\mathbf{R}_m^{(s)}\right\}\mathbf{W}^{(s)}\right]$$

subject to

$$\mathbf{W}^{(s)\top}\mathbf{W}^{(s)} = \mathbf{I}.$$

Therefore, $\mathbf{W}^{(s)}$ is updated by taking the leading $k_s$ eigenvectors of

$$\sum_{m=1}^{M}\mathbf{R}_m^{(s)\top}\mathbf{K}_{m,s}\mathbf{R}_m^{(s)}.$$

That is,

$$\boxed{\mathbf{W}^{(s)} = \operatorname{eigv}_{k_s}\left(\sum_{m=1}^{M}\mathbf{R}_m^{(s)\top}\mathbf{K}_{m,s}\mathbf{R}_m^{(s)}\right).} \tag{4}$$

**Update for $\mathbf{W}_m^{(\ell)}$.** To update the slice-specific loading matrix for slice $m$, define the residual after removing the shared component,

$$\mathbf{R}_m^{(\ell)} = \mathbf{Y}_m - \mathbf{Z}_m^{(s)}\mathbf{W}^{(s)\top}.$$

The slice-specific update solves

$$\min_{\mathbf{Z}_m^{(\ell)},\mathbf{W}_m^{(\ell)}} \left\|\mathbf{R}_m^{(\ell)} - \mathbf{Z}_m^{(\ell)}\mathbf{W}_m^{(\ell)\top}\right\|_F^2 + \lambda_\ell \operatorname{tr}\left(\mathbf{Z}_m^{(\ell)\top}\mathbf{L}_m\mathbf{Z}_m^{(\ell)}\right),$$

subject to

$$\mathbf{W}_m^{(\ell)\top}\mathbf{W}_m^{(\ell)} = \mathbf{I}, \qquad \mathbf{W}_m^{(\ell)\top}\mathbf{W}^{(s)} = \mathbf{0}.$$

For fixed $\mathbf{W}_m^{(\ell)}$, the optimal score matrix is

$$\mathbf{Z}_m^{(\ell)} = \mathbf{K}_{m,\ell}\mathbf{R}_m^{(\ell)}\mathbf{W}_m^{(\ell)}, \qquad \mathbf{K}_{m,\ell} = \left(\mathbf{I}_{n_m} + \lambda_\ell \mathbf{L}_m\right)^{-1}.$$

Substituting this into the objective reduces the problem to

$$\max_{\mathbf{W}_m^{(\ell)}} \operatorname{tr}\left[\mathbf{W}_m^{(\ell)\top}\mathbf{R}_m^{(\ell)\top}\mathbf{K}_{m,\ell}\mathbf{R}_m^{(\ell)}\mathbf{W}_m^{(\ell)}\right],$$

subject to

$$\mathbf{W}_m^{(\ell)\top}\mathbf{W}_m^{(\ell)} = \mathbf{I}, \qquad \mathbf{W}_m^{(\ell)\top}\mathbf{W}^{(s)} = \mathbf{0}.$$

Let

$$\mathbf{P}_s^{\perp} = \mathbf{I}_p - \mathbf{W}^{(s)}\mathbf{W}^{(s)\top}$$

denote the projection matrix onto the orthogonal complement of the shared loading space. Enforcing $\mathbf{W}_m^{(\ell)}$ to lie in this orthogonal complement gives the projected eigenproblem

$$\mathbf{P}_s^{\perp}\mathbf{R}_m^{(\ell)\top}\mathbf{K}_{m,\ell}\mathbf{R}_m^{(\ell)}\mathbf{P}_s^{\perp}.$$

Therefore,

$$\boxed{\mathbf{W}_m^{(\ell)} = \operatorname{eigv}_{k_\ell}\left(\mathbf{P}_s^{\perp}\mathbf{R}_m^{(\ell)\top}\mathbf{K}_{m,\ell}\mathbf{R}_m^{(\ell)}\mathbf{P}_s^{\perp}\right), \qquad \mathbf{P}_s^{\perp} = \mathbf{I}_p - \mathbf{W}^{(s)}\mathbf{W}^{(s)\top}.} \tag{5}$$

After obtaining the leading eigenvectors, $\mathbf{W}_m^{(\ell)}$ may be re-orthonormalized numerically to ensure

$$\mathbf{W}_m^{(\ell)\top}\mathbf{W}_m^{(\ell)} = \mathbf{I}, \qquad \mathbf{W}_m^{(\ell)\top}\mathbf{W}^{(s)} = \mathbf{0}.$$

**Summary of Updates.** The alternating minimization therefore consists of the following updates:

$$\mathbf{Z}_m^{(s)} = (\mathbf{I}_{n_m} + \lambda_s \mathbf{L}_m)^{-1} \mathbf{Y}_m \mathbf{W}^{(s)}, \tag{6}$$

$$\mathbf{Z}_m^{(\ell)} = (\mathbf{I}_{n_m} + \lambda_\ell \mathbf{L}_m)^{-1} \mathbf{Y}_m \mathbf{W}_m^{(\ell)}, \tag{7}$$

$$\mathbf{W}^{(s)} = \mathrm{eigv}_{k_s} \left( \sum_{m=1}^{M} \mathbf{R}_m^{(s)\top} \mathbf{K}_{m,s} \mathbf{R}_m^{(s)} \right), \tag{8}$$

$$\mathbf{W}_m^{(\ell)} = \mathrm{eigv}_{k_\ell} \left( \mathbf{P}_s^{\perp} \mathbf{R}_m^{(\ell)\top} \mathbf{K}_{m,\ell} \mathbf{R}_m^{(\ell)} \mathbf{P}_s^{\perp} \right), \tag{9}$$

where

$$\mathbf{R}_m^{(s)} = \mathbf{Y}_m - \mathbf{Z}_m^{(\ell)} \mathbf{W}_m^{(\ell)\top}, \qquad \mathbf{R}_m^{(\ell)} = \mathbf{Y}_m - \mathbf{Z}_m^{(s)} \mathbf{W}^{(s)\top},$$

and

$$\mathbf{K}_{m,s} = (\mathbf{I}_{n_m} + \lambda_s \mathbf{L}_m)^{-1}, \qquad \mathbf{K}_{m,\ell} = (\mathbf{I}_{n_m} + \lambda_\ell \mathbf{L}_m)^{-1}.$$

### S4 Replication-aware metafeatures for DDCIS and IBC slices

Figure 1 identifies the most important metafeatures shared across 28 slices used in Section 4.3.

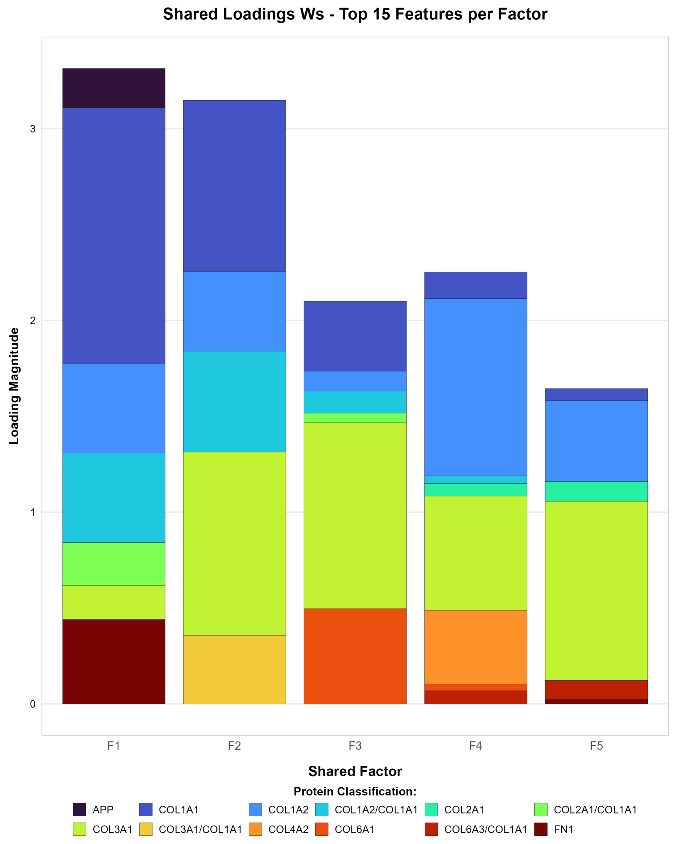


Figure 1: Reproducible metafeatures for DCIS and IBC slices in Section 4.3.
